# Human breast milk enhances intestinal mucosal barrier function and innate immunity in a pediatric human enteroid model

**DOI:** 10.1101/2021.02.17.431653

**Authors:** Gaelle Noel, Julie G. In, Jose M. Lemme-Dumit, Lauren R. DeVine, Robert N. Cole, Anthony L. Guerrerio, Olga Kovbasnjuk, Marcela F. Pasetti

**Affiliations:** Department of Pediatrics, Center for Vaccine Development and Global Health, University of Maryland School of Medicine, Baltimore, MD.; Department of Internal Medicine, Division of Gastroenterology and Hepatology, University of New Mexico Health Science Center, Albuquerque, NM.; Department of Medicine, Division of Gastroenterology and Hepatology, Johns Hopkins University School of Medicine, Baltimore, MD.; Department of Biological Chemistry, Johns Hopkins Mass Spectrometry and Proteomics Facility, Johns Hopkins University School of Medicine, Baltimore, MD.; Department of Pediatrics, Johns Hopkins University School of Medicine, Baltimore, MD.

## Abstract

Breastfeeding has been associated with long lasting health benefits. Nutrients and bioactive components of human breast milk promote cell growth, immune development, and shield the infant gut from insults and microbial threats. The molecular and cellular events involved in these processes are ill defined. We have established human pediatric enteroids and interrogated maternal milk’s impact on epithelial cell maturation and function in comparison with commercial infant formula. Colostrum applied apically to pediatric enteroid monolayers reduced ion permeability, stimulated epithelial cell differentiation, and enhanced tight junction function by upregulating occludin expression. Breast milk heightened the production of antimicrobial peptide α-defensin 5 by goblet and Paneth cells, and modulated cytokine production, which abolished apical release of pro-inflammatory GM-CSF. These attributes were not found in commercial infant formula. Epithelial cells exposed to breast milk elevated apical and intracellular pIgR expression and enabled maternal IgA translocation. Proteomic data revealed a breast milk-induced molecular pattern associated with tissue remodeling and homeostasis. Using a novel *ex vivo* pediatric enteroid model, we have identified cellular and molecular pathways involved in human milk-mediated improvement of human intestinal physiology and immunity.

## INTRODUCTION

The human gastrointestinal epithelium is a selective physical and chemical barrier that separates the luminal content from the serosal compartment and inner host tissues (1). It enables transport of electrolytes and nutrients, and provides a first line of defense against pathogens by engaging innate and adaptive mucosal immune components (2). The intestinal epithelium and associated mucosal immune environment progressively develop and mature from early fetal stages through childhood by means of genetic and external signals (3, 4). Human milk, rich in essential macronutrients, bioactive molecules (i.e., growth factors, antimicrobial peptides, complex oligosaccharides), and immune components including immunoglobulins, cytokines, and immune cells, supports tissue development and protects infants against infectious agents (5). Human milk is also a source of and helps establish a healthy microbiota in infants (6). Improvement of chronic and acute diseases (e.g., necrotizing enterocolitis, inflammatory bowel diseases, and intestinal and pulmonary infections) has been attributed to breastfeeding (7–10). Because of its countless benefits, breastfeeding has been recommended at least during the first 6 months of life (11). Current knowledge of the health-promoting benefits of human breast milk remains empiric or primarily descriptive, having been derived from observational or epidemiologic studies. The cellular and molecular mechanisms underlying the effects of maternal milk in the pediatric gut and physiologic pathways involved remain ill characterized. One of the reasons for this gap in knowledge is the lack of reliable models that could recapitulate the effect of human milk on the development and maintenance of a healthy pediatric human gut and its origin in modulating systemic effects. Studies using intestinal cancer cell lines including HT-29, T84, and Caco-2 cells or short-lived primary epithelial cells obtained from animals fail to reproduce the normal physiological responses of infant intestinal epithelium (12–15). Additionally, these immortalized cultures consist mainly of enterocytes and lack intestinal segment- and age-specificity needed for study of the complex multicellular and diverse composition of the human intestinal epithelium.

In this study, we described the establishment of an *ex vivo* pediatric human enteroid model derived from intestinal Lgr5^+^ stem cells and a mechanistic interrogation of the effects of human breast milk in the intestinal epithelium. Human intestinal enteroids (HIEs) recapitulate the crypt-villus cell axis and the segment-specific physiology (duodenum, jejunum, ileum) of the adult human small intestine (16, 17). Technical advantages of HIEs include their capacity for long-term growth (years), which preserves donor genotype, and forming polarized monolayers with easy access to apical and basolateral epithelial cell surfaces, which avoids the cumbersome manipulation of 3D structures (18). Herein, we present a side-by-side comparison of the molecular and cellular events affected by human milk vs. commercial infant formula in human pediatric enteroids. Outcome analyses included pediatric intestinal tissue morphology and maturation, ion and epithelial barrier permeability, antimicrobial and immune functions, and epithelial cell secretome.

## RESULTS

### Pediatric and adult enteroid monolayers exhibit distinct cell morphology and maturation features

To mechanistically interrogate the physiological effects of human breast milk in the pediatric gut, differentiated enteroid monolayers were established from duodenal biopsies of healthy 2-and 5-year-old children who underwent diagnostic endoscopy at The Johns Hopkins Hospital, using methods previously described (19, 20); these monolayers are hereafter referred to as 2PD and 5PD, respectively. The cell morphology, permeability and barrier integrity of the pediatric monolayers were compared with those derived from adult duodenal tissue. Differentiated (villus-like) enterocytes of pediatric origin were significantly shorter than their adult counterparts as revealed by confocal microscopy images (Figure 1A) and epithelial cell height measurement (Figure 1B). Analysis of the epithelial barrier function by transepithelial electrical resistance (TER) revealed increased paracellular ion permeability in the pediatric-as compared to the adult-derived monolayers (Figure 1C).

**Figure 1.**
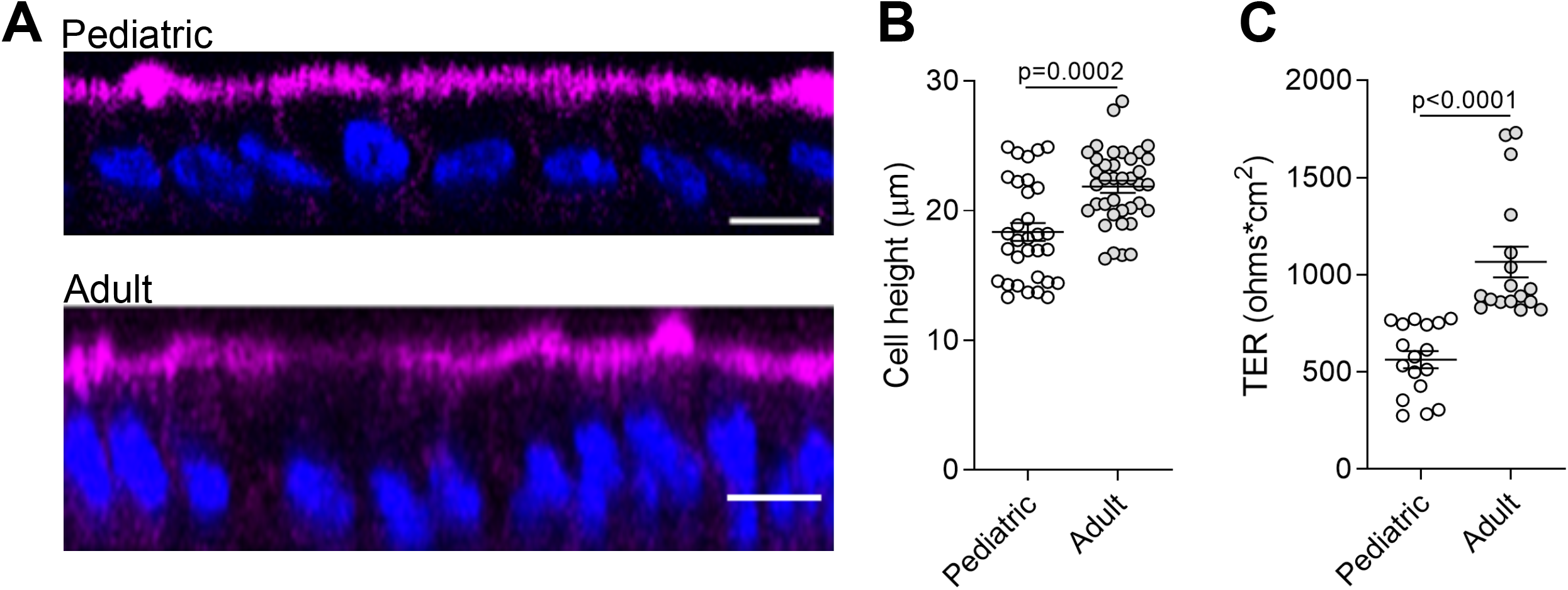
Pediatric and adult enteroid monolayers exhibit distinct maturation features. (**A**) Confocal microscopy images (XZ projections) depicting the difference in epithelial cell height between pediatric and adult enteroid monolayers. Actin, magenta; DNA, blue. Scale bar=20 μm. (**B**) Epithelial cell heights quantified by immunofluorescent confocal microscopy analysis (≥8 different view fields). (**C**) TER values of enteroid monolayers. Images are representative of three independent experiments (A). Data shown in (B) and (C) represent the mean ± SEM from three (B) or two (C) independent experiments that included *n*=8-12 enteroid monolayers/group per experiment. Each symbol represents an independent monolayer. (A-C) All measurements included 2 pediatric- and 3 adult-derived monolayers. (B, C) p-values were calculated by Student’s *t* test.

### Human breast milk improves pediatric epithelial barrier function

We next examined the effect of human breast milk (colostrum) on pediatric intestinal barrier function. Breast milk was applied to the apical side of differentiated pediatric enteroid monolayers, and TER values were monitored daily for 48h. Monolayers exposed to human breast milk exhibited higher TER values as compared to non-treated controls (Figures 2A and B). A dose-response effect was observed, with the 20% (v/v) treatment resulting in higher TER values as compared to 2% (v/v) (Figure 2A). This observation was consistent in multiple experiments using both lines; the 20% (v/v) solution was therefore selected for subsequent experiments. We next compared ion permeability of pediatric monolayers treated with human milk vs. commercial infant formula (also resuspended at 20% w/v). Human breast milk significantly and reliably increased TER levels in both 2PD and 5PD monolayers as compared to non-treated controls and remained elevated or further improved with prolonged exposure (Figures 2A and B). By contrast, ion permeability was modestly affected by infant formula; TER values increased only in the 5PD monolayer at 48h of treatment (Figure 2B, right panel). In addition to transepithelial ion permeability by TER, paracellular molecular permeability was examined by exposing breast milk- and infant formula-treated pediatric monolayers to FITC-labelled 4kDa dextran for up to 2h. No differences were observed in the amount of dextran recovered from the basolateral side regardless of treatment (data not shown) confirming integrity of the epithelial barrier.

**Figure 2.**
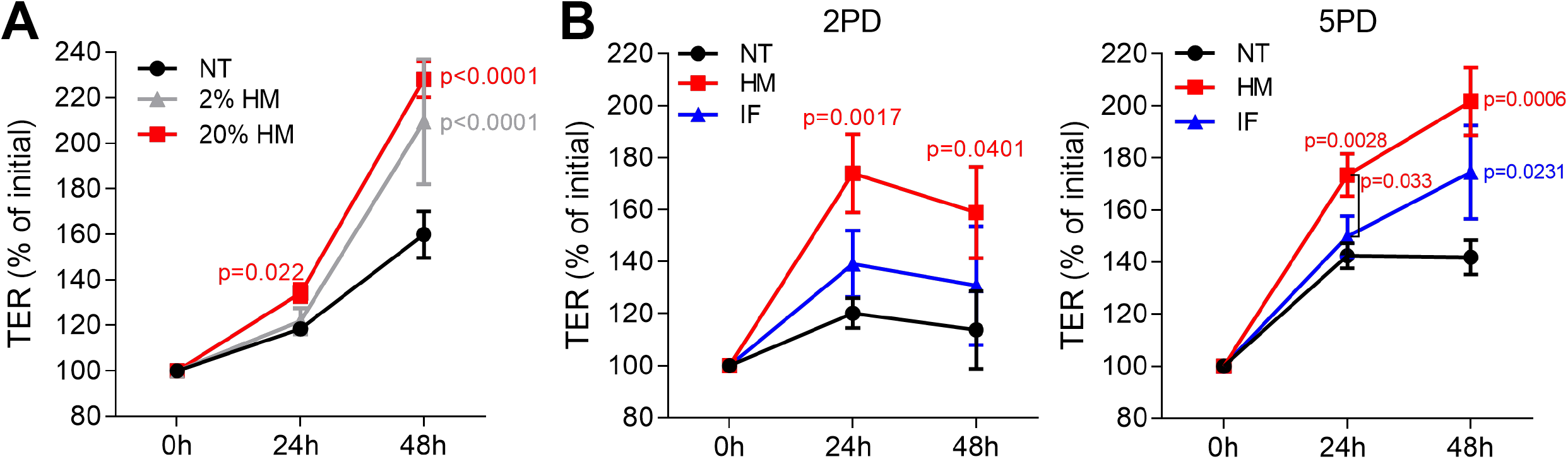
Human milk decreases ion permeability of the pediatric intestinal epithelium. (**A**) TER values of 2PD monolayers apically treated with 2% or 20% (v/v) of human milk (HM). (**B**) TER measurement of 2PD and 5PD monolayers apically treated with 20% (v/v) of HM or 20% (w/v) of commercial infant formula. Mean ± SEM. are shown. Data are representative of three independent experiments with *n*=3-6 enteroid monolayers/group per experiment. p-values were calculated by one-way-ANOVA with Šidák’s post-hoc analysis. Unless indicated, p-values correspond to treated vs. non-treated controls.

### Human breast milk increases the expression of the tight junction (TJ) protein occludin

Maternal milk enhancement of TER values prompted us to investigate its effect on expression of TJ proteins, which seal the paracellular space of the intestinal epithelia and regulate passage of ions and small molecules. Occludin, a transmembrane protein of the TJ complex was selected for this analysis as crucial marker of epithelial differentiation and barrier function (21). Immunofluorescent imaging revealed occludin on the cell perimeter of all monolayers, regardless of treatment (Figure 3A). Strikingly, pediatric monolayers exposed to human milk exhibited a distinctive pattern of apical and condensed cytoplasmic vesicular expression of occludin (Figure 3A) that markedly contrasted with the perimeter-only expression of monolayers treated with infant formula. Quantitative analysis of the fluorescence intensity by confocal imaging revealed superior occludin expression in both pediatric monolayers treated with human breast milk as compared with monolayers treated with infant formula or untreated controls (Figure 3B). Of the two enteroid lines, the 2PD was the higher and more consistent responder (Figure 3B). Infant formula increased occludin expression modestly and occasionally, not reaching significance above the non-treated controls (Figure 3B). The granular occludin expression pattern induced by breast milk was observed not only in absorptive enterocytes, visible by their prominent apical brush border, but also in cells lacking brush border, which are typically secretory epithelial cell lineages such as Paneth cells, goblet cells, and enteroendocrine cells (our HIE monolayers were not induced to express M cells). To identify the specific cell types producing occludin, breast milk-treated monolayers were co-stained to detect the presence of occludin as well as lysozyme, a marker for Paneth cells, trefoil factor 3 (TFF3), a marker for goblet cells, and chromogranin A, a marker for enteroendocrine cells. Occludin granular pattern co-localized with both lysozyme and TFF3, but not with chromogranin A marker (Figure 3C). These results indicate that breast milk elevates occludin expression not only at the TJ but also in the cytoplasm and apical membrane of absorptive enterocytes as well as in Paneth cells and goblet cells.

**Figure 3.**
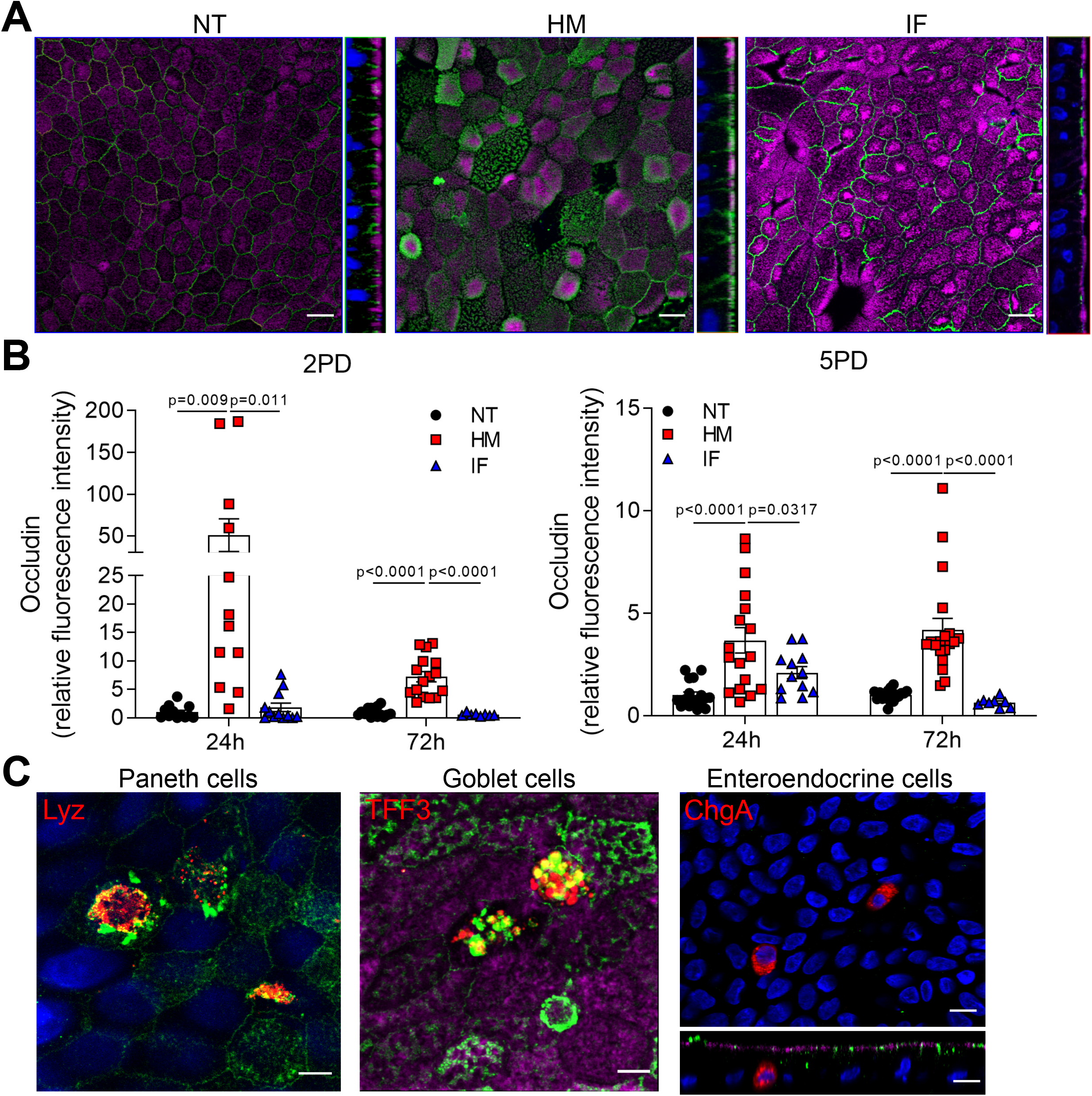
Human milk modulates occludin expression. (**A**) Confocal microscopy images (XY and YZ projections) of 2PD enteroid monolayers untreated (NT) or apically treated for 24h with HM (20%; v/v) or IF (20%; w/v). Occludin, green; actin, magenta. Scale bar=10 μm. (**B**) Relative fluorescence intensity of occludin quantified by confocal microscopy analysis of 2PD (left) and 5PD (right) monolayers treated with HM (20%; v/v) or IF (20%; w/v) for 24h and 72h. Mean ± SEM are shown. Data are pooled from three independents with *n*=4-6 enteroid monolayers/group per experiment. Each symbol indicates an independent monolayer. p-values were calculated by one-way-ANOVA with Šidák’s post-hoc analysis. (**C**) Confocal microscopy images (XY projections) of 5PD enteroid monolayers treated with HM for 48h. Occludin, green; lysozyme (Lyz; XY projection), red; trefoil factor 3 (TFF3; XY projection), red; chromogranin A (ChgA; XY and XZ projections), red; actin, magenta; DNA, blue. Paneth and goblet cells, scale bar=5 μm; enteroendocrine cells, scale bar=10 μm. (A and C) Data are representative of three independent experiments with *n*=3 enteroid monolayers/group per experiment.

### Human milk increases epithelial cell expression of innate immune mediators

The influence of breast milk on Paneth cell protein expression led us to examine its capacity to enhance Paneth cell function, and in particular the production of antimicrobial peptides such as Vα-defensin 5 (DEFA5), which helps maintain intestinal tolerance and homeostasis (22, 23). DEFA5 fluorescence intensity was greatly increased in breast milk-treated pediatric monolayers as compared to those treated with infant formula or non-treated controls (Figures 4A and B). Infant formula had no effect on DEFA5 expression. As expected, DEFA5 co-localized with lysozyme^+^ Paneth cells (Figure 4B). Surprisingly, a subpopulation of DEFA5-expressing cells that lacked the lysozyme marker was observed in human milk-treated monolayers (Figure 4B). Dual DEFA5^+^ and TFF3^+^ fluorescent staining revealed co-localization of these two markers, uncovering a breast milk-induced human goblet cell population with capacity to produce DEFA5 (Figure 4C).

**Figure 4.**
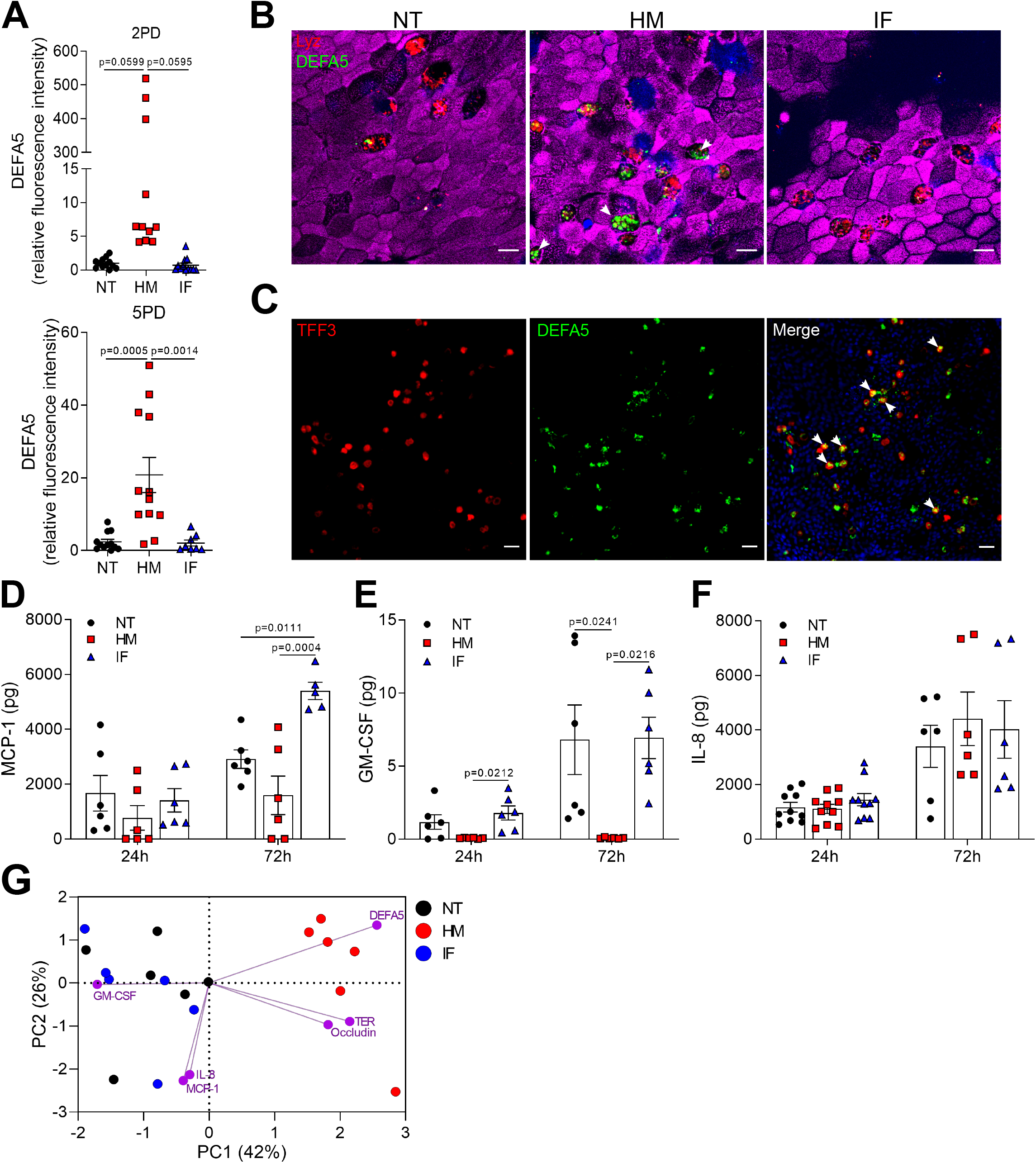
Human milk modulates epithelial innate immune mediators. (**A**) Relative fluorescence intensity of human DEFA5 quantified by confocal microscopy analysis of 2PD and 5PD monolayers NT or treated with HM (20%; v/v) or IF (20%; w/v) for 48h. (**B**) Representative confocal microscopy images (XY projections) of 5PD monolayer showing localization (arrowheads) of DEFA5 in Lyz-cells in HM-treated monolayer. DEFA5, green; Lyz, red; actin, magenta; DNA, blue. Scale bar=10 μm. (**C**) Representative confocal microscopy images (XY projections) of 5PD monolayer depicting co-localization (arrowheads) of TFF3 (red) and DEFA5 (green); DNA, blue. Scale bar=50 μm. (**D-F**) Total amount of MCP-1, GM-CSF, and IL-8 in the apical media of 2PD monolayer treated as described in (A) for 24h and 72h. (**G**) PCA plot from HM-, and IF-treated, and NT enteroid monolayers for 24h. PC, principal component. Variables analyzed: TER, occludin, DEFA5, MCP-1, GM-CSF, IL-8. (A, D-F). Mean ± SEM are shown. Data are representative of three independent experiments with *n*=6-12 enteroid monolayers/group per experiment. Each symbol indicates an independent monolayer. p-values were calculated by one-way-ANOVA with Tukey’s post-test for multiple comparisons.

We next examined the capacity of breast milk to modulate the production and secretion of cytokines and chemokines typically produced by intestinal epithelial cells. IL-10, IFN-γ, TNF-α, IL-6, IL-8, MCP-1, and GM-CSF were measured in the apical and basolateral milieu of treated and non-treated monolayers. IL-10 and IFN-γ in all conditions were below limit of detection (<0.7 pg). TNF-α and IL-6 were present at very low levels (<1 pg) and below the limit of detection in the non-treated controls, in both apical and basolateral compartments (data not shown). MCP-1, GM-CSF, and IL-8 were detected in apical media and for the most part, levels increased over time (Figures 4D-F). Treatment of pediatric monolayers with infant formula for 72h resulted in a marked increase of MCP-1 released apically as compared with non-treated monolayers. In contrast, a trend of reduced MCP-1 production was observed upon treatment with human milk (Figure 4D). GM-CSF was produced by untreated monolayers and by those treated with infant formula. In fact, infant formula produced a slight − yet not statistically significant − upregulation of GM-CSF at the 24h time point (Figure 4E). Conversely, apical GM-CSF secretion was abolished when monolayers were treated with human milk, at both time points tested (Figure 4E). Apical release of IL-8 remained unaffected by treatment (Figure 4F). Basolateral secretion of MCP-1, GM-CSF, and IL-8 was not influenced by treatment either (data not shown). A principal component analysis (PCA) was conducted combing 24h outcomes described above to visualize, in aggregate, the impact of breast milk and infant formula on epithelial cell physiology (the 24h time point was selected because it allowed for a complete dataset for all treatments). Monolayers untreated or exposed to infant formula clustered together and were largely distant from those exposed to breast milk by principal component 1 (Figure 4G). Breast milk treatment was associated with biomarkers of enhanced barrier function (DEFA5, occludin, and TER), whereas infant formula was linked to synthesis of pro-inflammatory cytokines (IL-8, MCP-1, and GM-CSF) (Figure 4G).

### Human milk sIgA translocates across pediatric enteroid monolayers

Breast milk contains a variety of immune mediators, including antibodies that shield immunologically naïve infants from health threats. Maternal immunoglobulins, in particular sIgA, support infant immune development and regulation, enacting long lasting benefits. Early colostrum has high levels of maternal sIgA and IgG, and hence our system enabled us to investigate their interaction with pediatric intestinal epithelial cells. Both 2PD and 5PD monolayers expressed secretory component (SC) of the polymeric immunoglobulin receptor (pIgR), which mediates IgA translocation across the intestinal epithelium as well as the neonatal Fc receptor (FcRn), responsible for transepithelial IgG transpot as shown by immunoblotting (Figures 5A and B). Confocal microscopy images revealed a diffuse cytoplasmic SC-pIgR expression in the non-treated controls, whereas epithelial cells exposed to breast milk exhibited not only intracellular but also basolateral and dense apical SC-pIgR expression (Figure 5A). Enhanced SC-pIgR expression in pediatric monolayers treated with breast milk was confirmed by immunoblotting (Figure 5C). Soluble SC-pIgR was detected in milk alone but not in infant formula (Figure 5C). We next compared apical to basolateral sIgA and IgG translocation in monolayers treated with breast milk vs. non-treated controls. Both sIgA and IgG were detected on basolateral side of milk-exposed epithelial cells; sIgA levels were significantly higher than those of the non-treated controls (Figure 5D).

**Figure 5.**
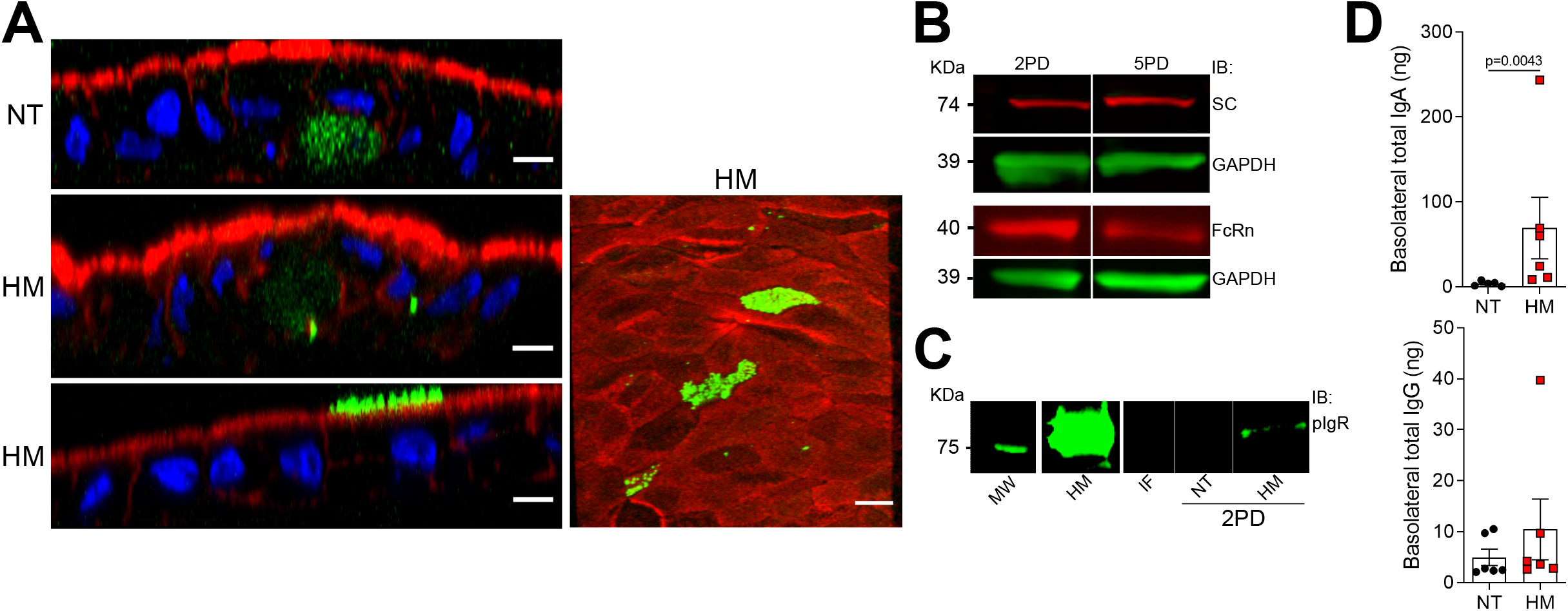
Breast milk enhances expression of pIgR and sIgA translocation across the epithelial monolayers. (**A**) Confocal microscopy images showing SC (left, XZ projections, scale bar=10 μm; right, XY projection, scale bar=5 μm) in 5PD enteroid monolayer NT or treated with 20% (v/v) of HM for 72h. SC, green; actin, red; DNA, blue. (**B**) Composite immunoblotting (IB) showing SC and FcRn expression in non-treated 2PD and 5PD monolayers. (**C**) IB showing pIgR expression in HM and IF, and 2PD monolayers NT or treated with 20% (v/v) HM. MW, molecular weight. (**D**) Total IgA and IgG in the basolateral media of pediatric monolayers treated for 48h with 20% (v/v) HM. Data represent mean ± SEM of three combined experiments, each including 2 monolayers/group per experiment. Each symbol indicates an independent monolayer. p-value was calculated by Student’s *t* test.

### Breast milk-induced protein upregulation and basolateral secretion by pediatric epithelial cells

The intestinal epithelium communicates with underlying tissues via secretion of nutrients, growth factors, cytokines, and regulatory peptides. Gut-derived molecules secreted to the basolateral compartment have the potential to disseminate systemically and act on remote tissues, exacting distant modulatory functions. To identify breast milk-induced molecules of intestinal origin that may have a wider (and possibly systemic) impact *in vivo*, we examined proteins secreted into the basolateral compartment of milk-exposed monolayers. Over 6000 proteins were identified by a proteomic analysis. To select the differentially expressed proteins from a total of 392 secreted proteins (with high false discovery rate), we applied the cutoffs: adjusted p-value ≤0.05 and log_2_ fold change at ±0.68. A total of 61 proteins had increased abundance in the breast milk-treated enteroids, whereas 21 were increased in the non-treated control (Figure 6A).

**Figure 6.**
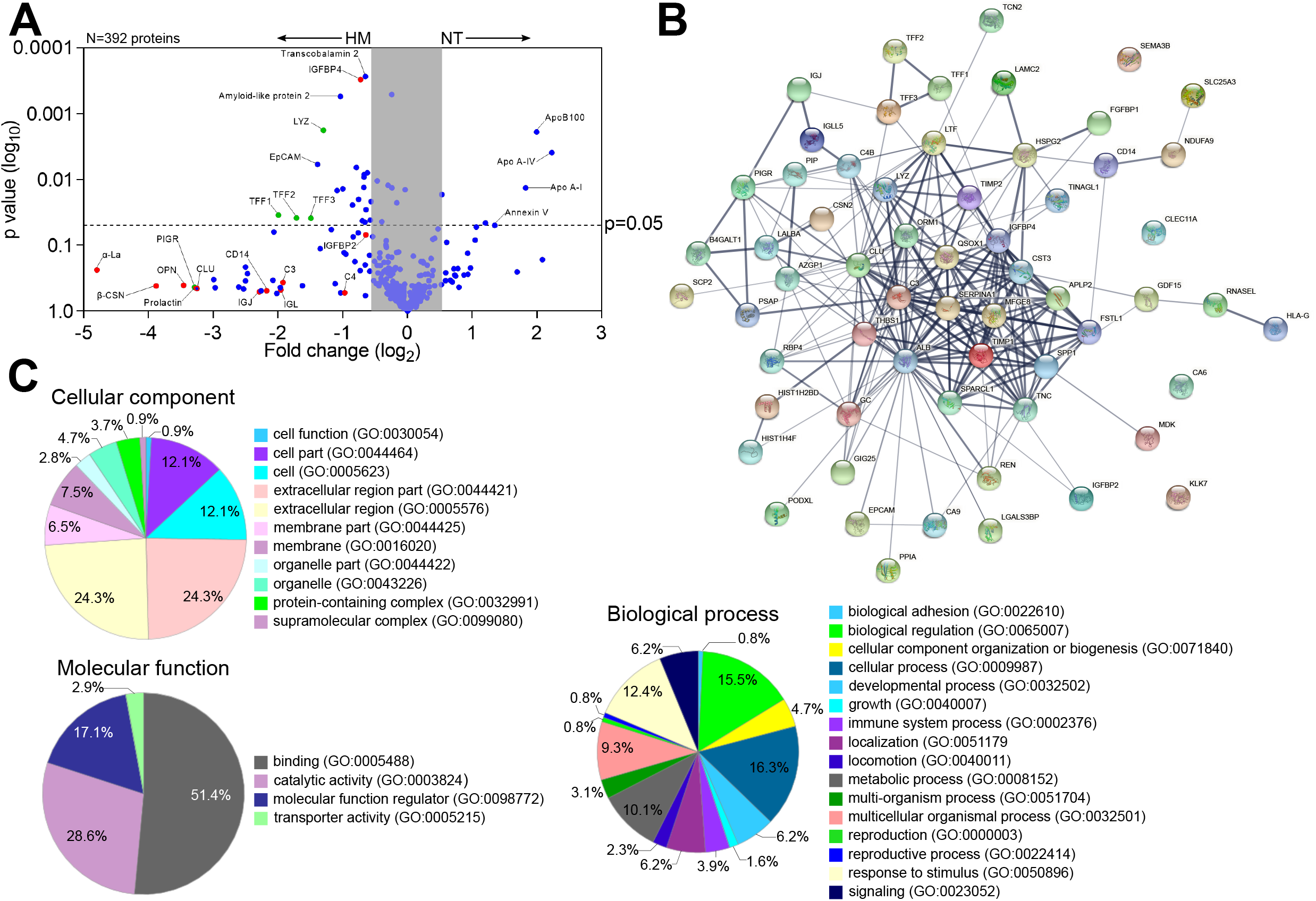
Human milk modifies epithelial cell protein expression and basolateral secretion. (**A**) Volcano plot of differential protein abundance (high false discovery rate) in the basolateral culture supernatant of 2PD monolayers NT (*n*=2) or treated with 20% HM (v/v) (*n*=3) for 24h. Red dots indicate HM unique proteins; blue dots indicate epithelial cell-derived proteins; green dots indicate proteins derived from both HM and epithelial cells. (**B**) Protein-protein interaction analysis of 61 upregulated proteins produced by HM-treated monolayers selected based on the cut off shown in (A). Medium confidence interaction score = 0.400. A thicker line between nodes indicates stronger protein-protein interaction. (**C**) Enrichment analysis of GO terms annotated for cellular component, molecular function, and biological process of the 61 upregulated proteins as described in (A).

Proteins derived from human milk were found in the basolateral compartment of breast milk-treated monolayers, indicating apical to basolateral transepithelial translocation. In addition, we observed increased levels of proteins related to mucosal protection and repair (e.g., TFF1-3, lysozyme C, amyloid-like protein), epithelial cell markers (e.g., EpCAM), growth factors (e.g., insulin-like growth factor-binding protein [IGFBP], fibroblast growth factor binding protein [FGFBP]), extracellular matrix remodeling proteins (e.g., metalloproteinase inhibitor proteins, basement membrane-specific heparan sulfate proteoglycan core protein) and cofactor carrier protein (e.g., transcobalamin 2) in the human breast milk-treated monolayers (Figure 6A). In contrast, the non-treated monolayers exhibited increased expression of the apolipoprotein family, and annexin V (Figure 6A). The interactions among proteins with increased abundance in the breast milk-treated enteroids were examined using the STRING v11.0 database. The analysis revealed a significant protein-protein interaction (p-value<1.0^-16^) among 57 of them (228 edges), whereas 4 proteins showed no interactions within the network (Figure 6B). These results indicate that most of the proteins secreted by milk-exposed enteroids do not act as independent entities but can deploy biological activity by either transient or stable association. A functional enrichment analysis was then performed utilizing the PANTHER and AMIGO2 classification database system to highlight the gene ontology (GO) terms annotated for cellular component, molecular function, and biological processes enriched within these protein sets (Figure 6C). The majority of the proteins were associated with the extracellular compartment (24.3%; GO:0044421, GO:0005576) as well as within the cell (12.1%; GO:0044464, GO:0005623) as constitutive protein with cytoplasmic or plasma membrane localization. The main molecular function identified was binding (51.4%; GO:0005488) followed by enzyme activity (28.6%; GO:0003824). In addition, these protein sets participate in multiple biological processes, including cell physiology, response to stimulus, metabolic functions, cell growth and maintenance, and immunity (Figure 6C).

## DISCUSSION

Human breast milk is a rich source of nutrients and bioactive components that promote infant growth and immune development. In this work, using an *ex vivo* pediatric intestinal stem cell-derived human enteroid model, we have identified distinct protein synthesized and cellular functions modulated by human breast milk. HIEs represent a cutting-edge technology that recapitulates the structural and functional features of the human gastrointestinal tissue. They have been used to interrogate gut physiology, host responses to microbes, drug activity, and cell-to-cell communication (24–29). A side-by-side comparison of pediatric-vs. adult-derived duodenal HIE monolayers revealed age-associated differences with the former exhibiting shorter columnar epithelial cells and reduced TER, consistent with a less mature epithelial cell phenotype. Reduced enterocyte height has been reported in duodenal biopsies of infants, as compared to adult subjects (30). Together, these results suggest that intestinal epithelial cell development continues through childhood. They also demonstrate that age-specific cell morphology is preserved in the HIEs.

Several unique molecular events associated with human milk improvement of pediatric intestinal health were observed. The first was the ability of breast milk (colostrum) to enhance epithelial barrier function by reducing ion permeability and upregulating expression of the TJ complex regulator occludin. The breast milk-treated monolayers exhibited an unusual pattern of upregulated occludin protein expression. Occludin was detected not only at the (expected) intercellular junctions but also on the apical plasma membranes of absorptive enterocytes as well as Paneth and goblet cells. Condensed occludin-containing vesicles were spread intracellularly. Apical occludin localization has been reported recently in mouse organoids, primarily in intestinal stem cells and Paneth cells, and less abundantly in enterocytes and goblet cells, and its presence associated with reduced paracellular permeability (31). A regulatory mechanism that involves recruitment of occludin contained in cytoplasmic vesicles or in the apical plasma membrane (via differential phosphorylation) for TJ formation has been proposed (32); under this model, the extra junctional localization may represent protein reservoirs that enable prompt TJ formation required by dynamic metabolic and physiological processes. To the best of our knowledge, this is the first demonstration of apical and cytoplasmic multi-lamellar occludin expression by human pediatric intestinal cells upregulated in response to breast milk.

A second key observation was the capacity of human milk to substantially increase production of human DEFA5, a peptide that contributes to innate host defense against enteropathogens and promotes intestinal homeostasis by limiting inflammation and microbial translocation (22, 33, 34). DEFA5 was produced not only by Paneth cells (the typical producers of antimicrobial molecules) but also by mucus-producing goblet cells. Production of DEFA5 by intestinal villous TFF3^+^ (goblet cells) but not lysozyme^+^ cells has been documented in human ileal biopsies (35). Goblet and Paneth cells derive from a common secretory cell progenitor under the regulation of ETS transcription factor Spdef (36). Lgr5^+^ stem cells and Paneth cells are abundant in crypt-like, non-differentiated HIEs. The lifespan of Paneth cells in enteroids is approximately 30 days, regardless of differentiation, as was previously shown in adult differentiated 3D enteroids (16). By contrast, the expression of DEFA5 in TFF3^+^ goblet cells, which mark the differentiated small intestinal epithelium, is a new finding and may reflect a differentiating cell lineage stage prompted by breast milk-derived growth factors. The heightening production of TJ proteins and antimicrobial products induced by breast milk (but not infant formula) is consistent with the reported improved epithelial barrier of infants fed with breast milk over those fed by formula as determined by reduced ratio of lactulose-to-mannitol in urine (37).

A third important observation was the immune modulation associated with human milk treatment of pediatric epithelial cells. While infant formula increased the production of pro-inflammatory cytokines MCP-1 and GM-CSF, breast milk reduced MCP-1 levels and totally suppressed apical release of GM-CSF. Gut inflammatory diseases such as intestinal bowel disease and celiac disease coincide with elevated MCP-1 and GM-CSF in duodenal biopsies (38).

IL-8, an epithelial cell-derived neutrophil chemoattractant was produced by the pediatric intestinal epithelium. Although not overtly affected by treatment, IL-8 was associated with exposure to infant formula as shown by PCA analysis of early time-point outcomes. These results are consistent with the anti-inflammatory properties of human milk, which, in the pediatric gut are deployed by reducing or abolishing steady state levels of signals that may activate or recruit phagocytic cells and enhance pro-inflammatory cytokines (i.e., GM-CSF and MCP-1) (39).

Different from adult HIEs, the pediatric HIEs did not produce substantial levels of TGF-β1, IFN-γ, IL-6, and TNF-α (19); these findings suggest that beyond the immune modulation of maternal milk, the pediatric intestinal epithelium is intrinsically programmed to silence signals that trigger inflammatory processes.

Human milk’s composition is complex and dynamic, and encompasses a vast diversity of soluble components that act as prebiotics, antiadhesives, antimicrobials, as well as molecules that affect cellular physiology, shield the host from inflammatory and pathogenic insults (40), Vand promote healthy gut development. Bioactive components with attributed anti-inflammatory and homeostatic function in human milk include IL-10, TGF-β, antioxidants, and enzymes such as lysozyme, glutathione peroxidase, and catalase (41). Additionally, human milk provides a variety of growth factors and tissue development/remodeling agents (42); proteomic analyses of human breast milk have been reported elsewhere (43, 44). We showed herein that many of these milk-derived components gain access to the subcellular space (see below). The exact molecules that trigger the effects described above and operatives, whether they work alone or in a synergistic/complementary manner, remain to be elucidated.

Maternal milk-derived sIgA provides an additional protective immune layer that excludes, neutralizes, and prevents microbial attachment to host cells (45). Mucosal dimeric IgA binds to pIgR on the basolateral surface of the epithelial cell membrane, is transported intracellularly and released at the apical surface, carrying a small portion of the pIgR-binding domain (46), the SC. Similar mechanism allows for IgM epithelial transport, whereas IgG employs the FcRn to bidirectionally cross epithelial tissues (47). Maternal antibodies provide antigen-specific defenses, support homeostasis, and promote infant immune development. In animal models, breast milk sIgA conferred long lasting benefits that included maintenance of a healthy microbiota and regulation of epithelial cell gene expression (48). A fourth relevant finding was the visualization of pIgR in the apical and basolateral membrane of breast milk-treated enterocytes. Breast milk itself contained an abundance of soluble SC-pIgR, but none was detected in commercial infant formula. The soluble SC-pIgR in maternal milk likely originates from maternal cellular debris. Free SC in human milk can bind enteric pathogens and toxins, and thus boosts non-specific host defenses (49) in the infant gut. We detected apical-to-basal sIgA transport in the maternal milk-exposed pediatric monolayers. This process supports intracellular pathogen neutralization and delivery of luminal antigens to lamina propria dendritic cells to induce tolerance or subepithelial phagocytic cells to imprint antigen specific immunity (50). FcRn detection in the pediatric tissue confirms expression of this receptor beyond infancy. Others have reported FcRn being expressed in human intestinal epithelial cells (51, 52).

We were unable to detect translocation of maternal IgG, despite this process being documented in animal models and cell lines (53). The variable localization of FcRn and pH requirements may restrict apical-to-basolateral transport while basolateral-to-apical appears to be more prevalent (54). Studies of FcRn distribution, IgG interaction and IgG immune complex translocation in pediatric HIEs are ongoing.

Beyond promoting a healthy gut, multiple and far reaching benefits have been attributed to human milk, including prevention of respiratory diseases, immune fitness, cognitive capacity, and overall physiological well-being (55) that endure into adolescence. Breast milk products released to the basal side of the epithelium could, conceivably, distribute systemically and thereby mediate long distant effects. Our proteomic analysis of breast milk-treated monolayers revealed a variety of molecules, some unique to breast milk, such as α-lactalbumin, β-casein, and prolactin, which had evidently translocated across the monolayers, and others that were produced by the milk-exposed pediatric intestinal cells. For the latter, a complex network of interacting biomolecules was revealed, with diverse functions including those affecting growth factors, immune and antimicrobial activity, tissue structure, and homeostasis, which confirms the broad and pleotropic nature of the processes affected by breast milk. The epithelial translocation of milk-derived proteins might have been facilitated by endocytosis of intact (undigested) molecules in our model. These proteins have health benefits by themselves. Milk α-lactalbumin, for example, shields soluble CD14 (sCD14) from proteolytic degradation (56), and sCD14 can bind lipopolysaccharide (LPS) and prevent inflammation and injury caused by soluble LPS or LPS-bearing organisms. β-casein is an immune modulator that regulates cell recruitment, ameliorates inflammation, and stimulates mucus production (57). Prolactin is a pleiotropic hormone that stimulates production of maternal milk. Expected benefits for the infant, based on animal studies, include reduction of anxiety and stress and neurogenesis (58). In addition, osteopontin prevents inflammation and epithelial damage in mouse DSS-colitis model (59).

A variety of breast milk-upregulated tissue-derived proteins were identified, including the TFF family, which maintains and restores gut mucosal homeostasis and regulates complement activation via decay-accelerating factor, DAF (60); the amyloid-like protein, which participates in intestinal metabolic processes and modulates expression of MHC class I molecules (61, 62); and, insulin growth factor binding protein, fibroblast growth factor, basement membrane-specific heparan sulfate protein, and metalloproteinase inhibitor – all of which contribute to epithelial cell growth, tissue remodeling, and barrier integrity (63–65). Other secreted proteins included transcobalamin 2, which facilitates the transport of vitamin B12 within the organs (66) and epithelial cell adhesion molecule (EpCAM), which localizes in the basal cell membrane and facilitates cell-to-cell interaction and proliferation (67). Complement proteins (C3 and C4) were also present in the basal media from breast milk-treated enteroids; C4 participates in complement activation via the classical and lectin pathway, whereas C3 is a converging substrate for all activating pathways; C3 cleavage into C3a and C3b, along with C5 cleavage, trigger the rest of the complement cascade. C3, C4, and other complement components are present in human breast milk (43, 44). Likewise, human intestinal epithelial cells produce complement proteins (68, 69). The origin of the complement proteins we identified is unclear. We surmise they derive from breast milk because synthesis of complement proteins by the intestinal epithelial cells reportedly requires pro-inflammatory signals (downregulated by breast milk in our system) (70). Nonetheless, the fact that maternal complement molecules would trespass the pediatric epithelium is intriguing. Regardless of their source, complement can boost infant mucosal protective mechanisms (71).

Bovine colostrum has been shown to influence the proteome of HT-29 cells as well as epithelial cell glycosylation (72). We show, for the first time, that human milk influences the synthesis of multiple mediators of metabolic and physiologic functions that act locally or systemically. In summary, using a novel *ex vivo* pediatric HIE, several mechanisms associated with breast milk were identified that improve intestinal health: 1) cell differentiation and strengthening of the pediatric intestinal barrier by reduction of permeability and upregulation of TJ occludin with a unique expression pattern; 2) boosting of innate immunity by enhancing production of antimicrobial DEFA5 by Paneth and goblet cells; 3) immune modulation and passive immunization by increased production of pIgR and translocation of luminal sIgA; 4) reduction of pro-inflammatory cytokines; 5) translocation of breast milk proteins with anti-inflammatory and anti-microbial properties; and 6) expression of proteins responsible for tissue remodeling and mucosal homeostasis.

## Methods

### Study approval

Protocols for recruitment of human participants, obtaining informed consent, collecting and de-identifying biopsy samples were approved by the Johns Hopkins University School of Medicine (JHU SOM) Institutional Review Board (IRB) NA 00038329. Procedures for recruitment of mothers around delivery, obtaining informed consent and collection and de-identification of breast milk were approved under University of Maryland School of Medicine IRB HP-00065842.

### Generation of enteroid monolayers

Duodenal biopsies were obtained from 5 healthy individuals, two pediatrics (ages 2, 5) and three adults (ages 25, 27, and 81 years) through endoscopy or surgical procedure. Enteroids were generated from Lgr5^+^ intestinal crypts embedded in Matrigel (Corning, USA) in 24-well plates, as previously described (73). Enteroids were expanded in growth factor-enriched media containing Wnt3A, Rspo-1, and Noggin (18, 19). Multiple enteroid cultures were harvested with Culturex Organoid Harvesting Solution (Trevigen, USA), fragmented and re-suspended in expansion media and seeded (100 μl) on the inner surface of 0.4 μm Transwell inserts (Corning, USA), pre-coated with human collagen IV (Sigma-Aldrich, USA), and 600 μl of expansion media was added to the receiver plate well. Media was replenished every other day (20). Enteroid monolayer confluency was monitored by measuring TER, as previously described (20). Upon reaching confluency, monolayers were differentiated in media (DFM) free of Wnt3A and Rspo-1 for 5 days (20). All cultures were maintained at 37°C and 5% CO_2_.

### Breast milk preparation and monolayer treatment

Human colostrum was obtained from women 0-3 days post-delivery. Commercial infant formula powder (Similac® Advance® Abbot Nutrition) was resuspended in sterile distilled water following manufacturer’s instructions. Both human breast milk and infant formula suspensions were centrifuged twice (10 min each) at 3,000g. The soluble fractions were extracted, aliquoted, and stored at −80°C until use. Enteroid monolayers were treated apically with 100 μl of human milk or infant formula diluted 2 or 20% in DFM. Non-treated controls were treated with 100 μl of DFM. TER was monitored daily while conducting experiments to ensure monolayer integrity.

### Dextran permeability assay

FITC-labelled 4 kDa dextran (Millipore Sigma, St. Louis, MO; 0.01% w/v in DFM) was added to the apical side of enteroid monolayers pre-treated with 20% of human milk or infant formula. Regular DFM (600 μl) was added to the basolateral side. Basolateral media (100 µl) was sampled at 30 min, 1 and 2h, and FITC-dextran content was measured by fluorescence intensity using an EnVision Multilabel Plate Reader (PerkinElmer, Waltham, MA). Sampled volume was replenished with fresh DFM.

### Immunofluorescence staining and confocal imaging

Enteroid monolayers were fixed for 40 min in 4% paraformaldehyde (Electron Microscopy Sciences, USA), washed with PBS for 10 min, permeabilized and blocked for 1h with PBS containing 15% fetal bovine serum, 2% BSA, and 0.1% saponin, all at room temperature (RT). After washing with PBS, monolayers were incubated overnight at 4°C with primary antibodies (diluted 1:100 in PBS). The following primary antibodies (Ab) were used: occludin (mouse monoclonal [mAb], clone OC-3F10, Thermo Fisher Scientific), TFF3 (rabbit polyclonal [pAb], Millipore Sigma), lysozyme EC 3.2.1.17 (rabbit pAb, Dako), DEFA5 (mouse mAb, clone 8C8, Millipore Sigma), and SC-166 (mouse mAb provided by Dr. A. Hubbard, Johns Hopkins University School of Medicine). Stained monolayers were washed with PBS (3 times, 10 min each) and incubated with secondary antibodies (diluted 1:100 in PBS) for 1h at RT. Secondary antibodies included goat anti-mouse Alexa Fluor-488 or -568, and goat anti-rabbit Alexa Fluor-488 or -568 (all Thermo Fisher Scientific). F-actin was detected by phalloidin Alexa Fluor-633, -647, or -568 (1:100; Thermo Fisher Scientific). Hoechst for nuclear/DNA labeling (Thermo Fisher Scientific) was used diluted 1:1000 in PBS. After incubation, cells were washed as described above, and mounted in FluorSave reagent (Millipore Sigma). Confocal images were taken using an LSM-510 META laser scanning confocal microscope (Zeiss, Germany) and ZEN 2012 imaging software (Zeiss) or BZ-X700 fluorescence microscope (Keyence, Japan) available through the Fluorescence Imaging Core of the Hopkins Basic Research Digestive Disease Development Center. For qualitative analysis, image settings were adjusted to optimize the signal. For quantitative analysis, the same settings were used across the samples, and protein-of-interest average intensity fluorescence was analyzed using MetaMorph software (Molecular Devices, CA).

### Protein extraction, immunoblotting, and proteomic analysis

Enteroid monolayers were lysed in cold lysis buffer (60 mM HEPES pH 7.4, 150 mM KCl, 5 mM Na3EDTA, 5 mM EGTA, 1 mM Na3VO4, 50 mM NaF, 2% SDS) supplemented with 1:100 of protease inhibitor cocktail (P8340, Millipore Sigma). Lysis buffer was applied to the apical surface, and cells were scraped and sonicated on ice (3 times at 10 sec pulses each time using 30% energy input). The lysates were centrifuged 10 min at 14000 rpm at 4°C, and the supernatant containing soluble and membrane proteins was collected. Total protein concentration was determined using a DC protein assay (Bio-Rad, CA). Proteins were separated on Novex Wedgewell 4-20% gradient Tris-glycine gels (Life Technologies, CA) and transferred to nitrocellulose membranes. The following primary antibodies were used for immunoblotting: polyclonal rabbit anti-pIgR (Abcam), monoclonal mouse anti-SC-166, and polyclonal rabbit anti-FcRn (Novus Biologicals) – all at a 1:250 dilution, and mouse monoclonal anti-GAPDH (clone 6C5, Abcam) at 1:1000 dilution. Secondary antibodies included goat anti-mouse Alexa Fluor-488 or -568 and goat anti-rabbit Alexa Fluor-488 or -568 (Thermo Fisher Scientific). Western blots were processed using the iBind Flex device (Life Technologies, Carlsbad, CA) and then imaged on an Odyssey CLx imager (LI-COR, Lincoln, NE). Proteomic analysis was conducted on basolateral media from pediatric monolayers treated with human milk (*n*=3) and non-treated control (*n*=2) through the Mass Spectrometry and Proteomics Facility, Johns Hopkins University School of Medicine.

### Cytokines/chemokines

Cytokines and chemokines were quantified using commercial electrochemiluminescence microarray kits (Meso Scale Diagnostic, Rockville, MD) following the manufacturer’s instructions. MCP-1, GM-CSF, and IL-8 levels were reported as the amount contained in the total volume of culture supernatant collected from the apical and basolateral side of the monolayers.

### Statistics

Statistical significances were calculated using the Student’s *t*-test to compare two groups, or one-way-ANOVA with Šidák’s or Tukey’s post-test as appropriate among more than two groups. PCA was performed by selecting PC with eigenvalues greater than 1.0 (Kaiser rule). Plots and statistical tests were performed using Prism software v9 (GraphPad, San Diego, CA). Differences were considered statistically significant at p-value ≤ 0.05.

## Author contributions

GN, JGI, and JML-D conducted experiments and analyzed data; JML-D compiled final figures; LD and RC conducted proteomics analysis; AG obtained pediatric biopsies; OK and MFP conceptualized the study, secured funding, designed experiments and data analysis. All authors contributed to the writing and editing of the manuscript.

## Acknowledgements

This work was supported by a Grand Challenge Exploration award (Bill and Melinda Gates Foundation) OPP 1118529 and in part, by NIH grants R01AI117734 (to MFP), P01 AI125181 (to MFP and OK) and K01 DK106323 (JGI). The authors acknowledge the Integrated Physiology and Imaging Cores of the Hopkins Digestive Disease Basic and Translational Research Core Center (P30 DK089502) and the Johns Hopkins Mass Spectrometry and Proteomics Core.

## Notes

### Competing Interest Statement

The authors have declared no competing interest.

### Summary of Updates

Discussion section updated.

## REFERENCES

1. Zihni C, et al. Tight junctions: from simple barriers to multifunctional molecular gates. Nat Rev Mol Cell Biol. 2016;17(9):564–80.

2. Peterson LW, and Artis D. Intestinal epithelial cells: regulators of barrier function and immune homeostasis. Nat Rev Immunol. 2014;14(3):141–53.

3. Torow N, et al. Neonatal mucosal immunology. Mucosal Immunol. 2017;10(1):5–17.

4. Stras SF, et al. Maturation of the Human Intestinal Immune System Occurs Early in Fetal Development. Dev Cell. 2019;51(3):357–73 e5.

5. Turfkruyer M, and Verhasselt V. Breast milk and its impact on maturation of the neonatal immune system. Curr Opin Infect Dis. 2015;28(3):199–206.

6. Ballard O, and Morrow AL. Human milk composition: nutrients and bioactive factors. Pediatr Clin North Am. 2013;60(1):49–74.

7. Bode L. Human Milk Oligosaccharides in the Prevention of Necrotizing Enterocolitis: A Journey From in vitro and in vivo Models to Mother-Infant Cohort Studies. Front Pediatr. 2018;6:385.

8. Jantscher-Krenn E, et al. The human milk oligosaccharide disialyllacto-N-tetraose prevents necrotising enterocolitis in neonatal rats. Gut. 2012;61(10):1417–25.

9. Barclay AR, et al. Systematic review: the role of breastfeeding in the development of pediatric inflammatory bowel disease. J Pediatr. 2009;155(3):421–6.

10. Oddy WH. Breastfeeding, Childhood Asthma, and Allergic Disease. Ann Nutr Metab. 2017;70 Suppl 2:26–36.

11. World Health Organization. WHO Recommendations on Postnatal Care of the Mother and Newborn. Geneva; 2013.

12. Sun D, et al. Comparison of human duodenum and Caco-2 gene expression profiles for 12,000 gene sequences tags and correlation with permeability of 26 drugs. Pharm Res. 2002;19(10):1400–16.

13. Drummond CG, et al. Enteroviruses infect human enteroids and induce antiviral signaling in a cell lineage-specific manner. Proc Natl Acad Sci U S A. 2017;114(7):1672–7.

14. Lin S, et al. Comparison of the transcriptional landscapes between human and mouse tissues. Proc Natl Acad Sci U S A. 2014;111(48):17224–9.

15. Pulendran B, and Davis MM. The science and medicine of human immunology. Science. 2020;369(6511).

16. Sato T, et al. Single Lgr5 stem cells build crypt-villus structures in vitro without a mesenchymal niche. Nature. 2009;459(7244):262–5.

17. Zachos NC, et al. Human Enteroids/Colonoids and Intestinal Organoids Functionally Recapitulate Normal Intestinal Physiology and Pathophysiology. J Biol Chem. 2016;291(8):3759–66.

18. In JG, et al. Human colonoid monolayers to study interactions between pathogens, commensals, and host intestinal epithelium. J Vis Exp. 2019(146).

19. Noel G, et al. A primary human macrophage-enteroid co-culture model to investigate mucosal gut physiology and host-pathogen interactions. Sci Rep. 2017;7:45270.

20. Staab JF, et al. Co-Culture System of Human Enteroids/Colonoids with Innate Immune Cells. Curr Protoc Immunol. 2020;131(1):e113.

21. Al-Sadi R, et al. Occludin regulates macromolecule flux across the intestinal epithelial tight junction barrier. Am J Physiol Gastrointest Liver Physiol. 2011;300(6):G1054–64.

22. Bevins CL, and Salzman NH. Paneth cells, antimicrobial peptides and maintenance of intestinal homeostasis. Nat Rev Microbiol. 2011;9(5):356–68.

23. Sankaran-Walters S, et al. Guardians of the Gut: Enteric Defensins. Front Microbiol. 2017;8:647.

24. In JG, et al. Epithelial WNT2B and Desert Hedgehog Are Necessary for Human Colonoid Regeneration after Bacterial Cytotoxin Injury. iScience. 2020;23(10):101618.

25. Liu L, et al. Mucus layer modeling of human colonoids during infection with enteroaggragative E. coli. Sci Rep. 2020;10(1):10533.

26. Co JY, et al. Controlling epithelial polarity: A human enteroid model for host-pathogen interactions. Cell Rep. 2019;26(9):2509-20.e4.

27. King AJ, et al. Inhibition of sodium/hydrogen exchanger 3 in the gastrointestinal tract by tenapanor reduces paracellular phosphate permeability. Sci Transl Med. 2018;10(456).

28. Lin SC, et al. Human norovirus exhibits strain-specific sensitivity to host interferon pathways in human intestinal enteroids. Proc Natl Acad Sci U S A. 2020;117(38):23782–93.

29. Chang-Graham AL, et al. Rotavirus induces intercellular calcium waves through ADP signaling. Science. 2020;370(6519).

30. Thompson FM, et al. Epithelial growth of the small intestine in human infants. J Pediatr Gastroenterol Nutr. 1998;26(5):506–12.

31. Pearce SC, et al. Marked differences in tight junction composition and macromolecular permeability among different intestinal cell types. BMC Biol. 2018;16(1):19.

32. Wong V. Phosphorylation of occludin correlates with occludin localization and function at the tight junction. Am J Physiol. 1997;273(6):C1859–67.

33. Salzman NH, et al. Enteric defensins are essential regulators of intestinal microbial ecology. Nat Immunol. 2010;11(1):76–83.

34. Ehmann D, et al. Paneth cell alpha-defensins HD-5 and HD-6 display differential degradation into active antimicrobial fragments. Proc Natl Acad Sci U S A. 2019;116(9):3746–51.

35. Cunliffe RN, et al. Human defensin 5 is stored in precursor form in normal Paneth cells and is expressed by some villous epithelial cells and by metaplastic Paneth cells in the colon in inflammatory bowel disease. Gut. 2001;48(2):176–85.

36. Gregorieff A, et al. The ets-domain transcription factor Spdef promotes maturation of goblet and paneth cells in the intestinal epithelium. Gastroenterology. 2009;137(4):1333–45 e1-3.

37. Catassi C, et al. Intestinal permeability changes during the first month: effect of natural versus artificial feeding. J Pediatr Gastroenterol Nutr. 1995;21(4):383–6.

38. Di Sabatino A, et al. Innate and adaptive immunity in self-reported nonceliac gluten sensitivity versus celiac disease. Dig Liver Dis. 2016;48(7):745–52.

39. Hamilton JA. GM-CSF in inflammation. J Exp Med. 2020;217(1).

40. Bode L. The functional biology of human milk oligosaccharides. Early Hum Dev. 2015;91(11):619–22.

41. Cacho NT, and Lawrence RM. Innate Immunity and Breast Milk. Front Immunol. 2017;8:584.

42. Ogra PL. Immunology of Human Milk and Lactation: Historical Overview. Nestle Nutr Inst Workshop Ser. 2020;94:11–26.

43. Zhu J, and Dingess KA. The Functional Power of the Human Milk Proteome. Nutrients. 2019;11(8).

44. Gao X, et al. Temporal changes in milk proteomes reveal developing milk functions. J Proteome Res. 2012;11(7):3897–907.

45. Cerutti A, and Rescigno M. The biology of intestinal immunoglobulin A responses. Immunity. 2008;28(6):740–50.

46. Brandtzaeg P. Secretory IgA: Designed for Anti-Microbial Defense. Front Immunol. 2013;4:222.

47. Pyzik M, et al. FcRn: The Architect Behind the Immune and Nonimmune Functions of IgG and Albumin. J Immunol. 2015;194(10):4595–603.

48. Rogier EW, et al. Secretory antibodies in breast milk promote long-term intestinal homeostasis by regulating the gut microbiota and host gene expression. Proc Natl Acad Sci U S A. 2014;111(8):3074–9.

49. Giugliano LG, et al. Free secretory component and lactoferrin of human milk inhibit the adhesion of enterotoxigenic Escherichia coli. J Med Microbiol. 1995;42(1):3–9.

50. Corthesy B. Multi-faceted functions of secretory IgA at mucosal surfaces. Front Immunol. 2013;4:185.

51. Israel EJ, et al. Expression of the neonatal Fc receptor, FcRn, on human intestinal epithelial cells. Immunology. 1997;92(1):69–74.

52. Latvala S, et al. Distribution of FcRn Across Species and Tissues. J Histochem Cytochem. 2017;65(6):321–33.

53. Dickinson BL, et al. Bidirectional FcRn-dependent IgG transport in a polarized human intestinal epithelial cell line. J Clin Invest. 1999;104(7):903–11.

54. Aaen KH, et al. The neonatal Fc receptor in mucosal immune regulation. Scand J Immunol. 2021;93(2):e13017.

55. Krol KM, and Grossmann T. Psychological effects of breastfeeding on children and mothers. Bundesgesundheitsblatt Gesundheitsforschung Gesundheitsschutz. 2018;61(8):977–85.

56. Spencer WJ, et al. Alpha-lactalbumin in human milk alters the proteolytic degradation of soluble CD14 by forming a complex. Pediatr Res. 2010;68(6):490–3.

57. Chatterton DE, et al. Anti-inflammatory mechanisms of bioactive milk proteins in the intestine of newborns. Int J Biochem Cell Biol. 2013;45(8):1730–47.

58. Torner L. Actions of Prolactin in the Brain: From Physiological Adaptations to Stress and Neurogenesis to Psychopathology. Front Endocrinol (Lausanne). 2016;7:25.

59. Woo SH, et al. Osteopontin Protects Colonic Mucosa from Dextran Sodium Sulfate-Induced Acute Colitis in Mice by Regulating Junctional Distribution of Occludin. Dig Dis Sci. 2019;64(2):421–31.

60. Andoh A, et al. Intestinal trefoil factor induces decay-accelerating factor expression and enhances the protective activities against complement activation in intestinal epithelial cells. J Immunol. 2001;167(7):3887–93.

61. Puig KL, et al. Amyloid precursor protein mediated changes in intestinal epithelial phenotype in vitro. PLoS One. 2015;10(3):e0119534.

62. Tuli A, et al. Amyloid precursor-like protein 2 increases the endocytosis, instability, and turnover of the H2-K(d) MHC class I molecule. J Immunol. 2008;181(3):1978–87.

63. Austin K, et al. IGF binding protein-4 is required for the growth effects of glucagon-like peptide-2 in murine intestine. Endocrinology. 2015;156(2):429–36.

64. Tassi E, et al. Impact of fibroblast growth factor-binding protein-1 expression on angiogenesis and wound healing. Am J Pathol. 2011;179(5):2220–32.

65. Cabral-Pacheco GA, et al. The Roles of Matrix Metalloproteinases and Their Inhibitors in Human Diseases. Int J Mol Sci. 2020;21(24).

66. Quadros EV, et al. Transcobalamin II synthesized in the intestinal villi facilitates transfer of cobalamin to the portal blood. Am J Physiol. 1999;277(1):G161–6.

67. Das B, et al. Enteroids expressing a disease-associated mutant of EpCAM are a model for congenital tufting enteropathy. Am J Physiol Gastrointest Liver Physiol. 2019;317(5):G580–G91.

68. Moon R, et al. Complement C3 production in human intestinal epithelial cells is regulated by interleukin 1beta and tumor necrosis factor alpha. Arch Surg. 1997;132(12):1289–93.

69. Kopp ZA, et al. Do antimicrobial peptides and complement collaborate in the intestinal mucosa? Front Immunol. 2015;6:17.

70. Andoh A, et al. Differential cytokine regulation of complement C3, C4, and factor B synthesis in human intestinal epithelial cell line, Caco-2. J Immunol. 1993;151(8):4239–47.

71. Ogundele M. Role and significance of the complement system in mucosal immunity: particular reference to the human breast milk complement. Immunol Cell Biol. 2001;79(1):1–10.

72. Morrin ST, et al. Interrogation of Milk-Driven Changes to the Proteome of Intestinal Epithelial Cells by Integrated Proteomics and Glycomics. J Agric Food Chem. 2019;67(7):1902–17.

73. Sato T, et al. Long-term expansion of epithelial organoids from human colon, adenoma, adenocarcinoma, and Barrett’s epithelium. Gastroenterology. 2011;141(5):1762–72.

